# Daytime heat exposure increases nighttime predation risk in a mangrove gastropod

**DOI:** 10.64898/2026.05.10.723115

**Authors:** Wissam A. Jawad, Rachel Collin, Christopher Dwane, Morgan W. Kelly

## Abstract

1. The frequency and intensity of heat events is increasing across marine and terrestrial ecosystems. Within the same ecological community, the relative exposure and sensitivity to heat stress may vary considerably among interacting species, like predators and prey. This can be especially true for species that interact at the aquatic-terrestrial interface, as well as for interactions between primarily nocturnal and diurnal species, making it difficult to predict how such communities will respond to habitat warming.
2. Thermal limit metrics such as CT_max_ are often assumed to equate with ‘ecological death’ because such temperatures impair behavioral activity and/or physiological functioning. Prey that are diurnally active can be more frequently exposed to temperatures that approach CT_max_ compared to their nocturnal predators, which may use thermal refuges during the day. Yet the impacts of daytime heat exposure on nighttime predation risk remain unknown.
3. Here, we compared the thermal environment, performance, and heat tolerance between the predatory blue crab, *Callinectus sapidus* and one of its prey species, the mangrove periwinkle *Littoraria anguilifera* in a tropical mangrove ecosystem. We examined how exposing prey to heat stress at and below their CT_max_ affected their capacity to avoid predation in the field at night when predation risk is highest.
4. We found that acute exposure to temperatures near CT_max_ during the day increased the prey species’ susceptibility to predation during recovery at night. Although both interacting predator and prey have high thermal tolerance, prey are exposed to conditions that already reach CT_max_, suggesting that current extremes in temperatures may already be influencing vulnerability to predation in this ecosystem.
5. Our results suggest that differential exposure to sublethal heat stress in diurnal prey relative to their predator, along with the subsequent impact of these exposures on predation risk, will play a role in shaping these interacting as climate warms.

## Introduction

Climate change is increasing mean global temperatures as well as the frequency, intensity and duration of extreme heat events (Fischer & Knutti 2015, Perkins et al. 2012). Extremes in temperatures are predicted to have disproportionate impacts on species survival due to acute exposure to physiologically stressful temperatures. For example, recent heat waves have been linked to mass die offs and population extirpations across marine (Leggat et al 2019, Smale & Wernberg 2013) and terrestrial taxa (Harvey et al. 2020, Ruthrof et al. 2018). However, not all species within a community are equally vulnerable. Variation in species responses to climate warming within the same ecological community can in part be attributed to asymmetries in species thermal sensitivity, as well as different levels of exposure to extreme conditions. For example, variation in thermal tolerance can predict responses to warming (Sorte et al. 2011), where species with lower heat tolerance can experience greater population declines as habitat temperatures increase compared to species with higher tolerance within the same habitat (Roeder et al. 2021). At the same time, species that occupy relatively cooler microhabitats and/or have a greater capacity to behaviorally thermoregulate can reduce their exposure to extreme temperatures (Huey et al. 2012, Apte et al. 2025), and thus may have a higher warming tolerance, defined as the difference between an organism’s upper thermal limit and maximum habitat temperatures (Deutch et al 2008, Pinsky et al. 2019, Jawad et al. 2026). Understanding the factors that contribute to interspecific variation in climate vulnerability will be vital for anticipating ecosystem responses to climate warming, especially between interacting species, where changes in ecological interactions can have cascading impacts on the ecosystem as a whole. Predator-prey interactions can profoundly influence ecosystem structure and diversity (Paine 1966, Power 1990, Estes et al. 2011) and the strength of such interactions is sensitive to temperature (Sanford 1999, Miller et al 2014, Uszko et al 2017). The thermal sensitivity of interacting species, and thus their responses to warming, can in part be predicted by their respective upper thermal tolerances relative to the current maximum temperatures in their environments (i.e. warming tolerance). In many ecosystems, however, the realized maximum temperatures experienced by organisms will depend on their relative mobility and use of microhabitats, which itself can be influenced by ecological pressures. For example, prey activity can be restricted to warmer microhabitats and times of the day compared to their predators because of fear-induced changes in behavior (Schmitz & Suttle 2001, Vaudo & Heithaus 2013, Jawad et al. 2024). This can lead to different realized thermal distributions between predators and prey, where predator presence can directly influence the habitat temperatures that prey experience. For example, increased risk of predation at night could induce prey to switch to diurnal foraging (Fraser et al. 2004, Hudgens et al. 2011), thereby reducing predation risk at night, but potentially increasing exposure to increased heat stress by being active during the day. Despite the ubiquity of diurnal activity in prey and nocturnal activities in predators, little is known about the impacts of increasing day time temperatures on prey vulnerability to nocturnal predation.

Differences in mobility and access to thermal refugia between predators and prey can also impact their relative evolved responses to heat stress (Menge and Sutherland 1987). Greater behavioral capacity to avoid extreme high temperatures in more mobile species could result in reduced selection for heat tolerance (Bogert 1949, Muñoz 2022). Prey with less mobility or restricted habitat use, on the other hand, may experience greater exposure to thermal stress and thus selection favoring increased physiological heat tolerance (McIntire and Miller 2025). Based on these attributes of predator-prey interactions, the trophic sensitivity hypothesis predicts that predators should have lower physiological tolerance to environmental stressors than their prey (Cheng et al 2017). In addition, previous work has demonstrated that prey have a lower and broader range of temperatures for behaviors associated with predator avoidance compared to predation behaviors (i.e. attack speeds), suggesting a ‘thermal life-dinner principle’ (Dell et al. 2011), such that the costs of reduced predator avoidance because of thermal sensitivity are greater (death) than predators missing a meal. Yet empirical tests comparing the thermal sensitivity and performance of predators and prey have yielded mixed findings, where in some contexts predators have the same or lower thermal performance than their prey (Cheng et al. 2017), while in other habitats, predators have higher thermal performance (Monaco et al. 2016) and higher thermal tolerance (Pintanel et al 2021) than their prey. Untangling the contexts and traits which may predict tolerance to warming in predators and prey thus remains a pressing question in predicting the impacts of climate warming on ecological communities.

Greater exposure to heat stress in prey could also impact their capacity to avoid predators as heat events become more frequent and intense. The critical thermal maximum (CT_max_) describes the temperature at which an organism experiences neuromuscular failure and loses locomotory control (Luttershmidt and Hutchison 1997, Ørsted et al. 2022). Generally, an implicit assumption is made that CT_max_ marks the point of ‘ecological death’, where organisms should experience severe ecological costs, such as greater susceptibility to predation because of reduced mobility. However, this has rarely been empirically demonstrated in predator-prey studies (but see Leong et al. 2023), and less still is understood about the effects of heat stress accumulated below CT_max_. Exposures to sublethal heat stress have the potential to reduce antipredator responses in prey in numerous ways, including the need for physiological recovery post-heat exposure (Joyce et al. 2022, Stillman et al 2025, Clements et al. 2025). For example, performance traits of marine gastropods can be reduced for several hours after exposure to acute heat stress (Hemraj et al. 2017). At the molecular level, heat shock protein activation can require heavy energy demands (Feder & Hofmann 1999), and genes associated with body maintenance can be expressed up to 24 hours after acute heat exposures in splash pool copepods (Vaidya et al. 2025), suggesting that organisms need to channel energy towards physiological recovery hours after heat stress is reduced. Such sustained responses would suggest that behavioral responses to ecological threats like predation could be reduced post heat exposure as heat events increase exposure to such conditions (Breedveld et al. 2025). Studies that pair physiological data with behavioral and functional trait responses to acute heat exposure will be important for estimating ecological vulnerability in prey (Stoks et al. 2017) and can aid in understanding the relationship between thermal limits and the impacts of stress accumulated below these limits (Rezende et al. 2014).

Intertidal gastropods, in particular tropical periwinkles (littorines), represent a notable example of extreme physiological adaptation to temperature stress (Marshall et al. 2011, Zhang et al. 2025). Littorines can physiologically withstand a wider range of temperatures than those at which they are behaviorally active in (Evans 1948, Monaco et al. 2017) and thus can exploit hot microhabitats in the upper intertidal zone, enabling them to escape subtidal predators such as portunid swimming crabs (Cannicci et al. 1996, Silliman & Bertness 2001, Jawad et al. 2024). Swimming crabs are largely subtidal and therefore do not experience the same thermal extremes as their high intertidal prey. Swimming crabs’ prey on littorines by extending their claws out of the water and capturing snails within reach, and are most active at night, further reducing the thermal stress experienced when feeding. Foraging at times of day when thermal stress is lower may be especially important for predators because attack potential in predators may be more negatively impacted by thermal stress compared to prey escape potential (Öhlund et al. 2015). Although thermal tolerance traits in littorines have received much attention (Wang et al. 2022, Marshall et al. 2015, Han et al. 2019), variation in thermal sensitivity and exposure to heat stress between littorines and their subtidal predators remain unknown, making it difficult to predict the relative responses of the interacting species, and thus to predict a potential for shifts in their interactions under future conditions. In addition, although many tropical littorines have a high thermal tolerance (Marshall et al. 2015), how sublethal heat stress impacts their capacity to avoid predation remains unknown.

Here, we examined the predation threat associated with microhabitat use, relative thermal sensitivity, and impacts of acute heat stress in the tropical mangrove periwinkle (*Littoraria anguilifera*, Lamarck 1822), alongside that of a key predator, the blue crab (*Callinectus sapidus*, Rathbun 1896) in a mangrove infringed bay in Colon Island in Bocas del Toro, Panama. *L. angulifera* is locally abundant on red mangroves (*Rhizophora mangle*, Linnaeus 1753), where it feeds on biofilms and algae near the water line on the prop roots during the day (Fig. 1), before migrating to the higher portions of the trees at night. Previous research has demonstrated that distance above the water line is negatively correlated with predation threat (Duncan and Szelistowski 1998, Rochette and Dill 2000) and can also tradeoff with behavioral thermoregulation (Jawad et al 2024). Littorines generally have wide apertures and lack defensive shell morphology, making anti-predator behavior crucial for snails in this genus (Vermeij 1979). *C. sapidus* are abundant in calm mangrove and estuary habitats, where they crawl and swim through mangrove roots and other vegetation to feed on prey including *L. angulifera*, which they consume by peeling or crushing the shell altogether. Handling and consumption of *L. angulifera* by *C. sapidus* can be rapid (<30 s for a single adult, unpublished data). Crabs commonly rest in reefs during the day, which may offer refuge from bird predators and heat in the shallower waters where they feed around the mangroves.

**Fig. 1.**
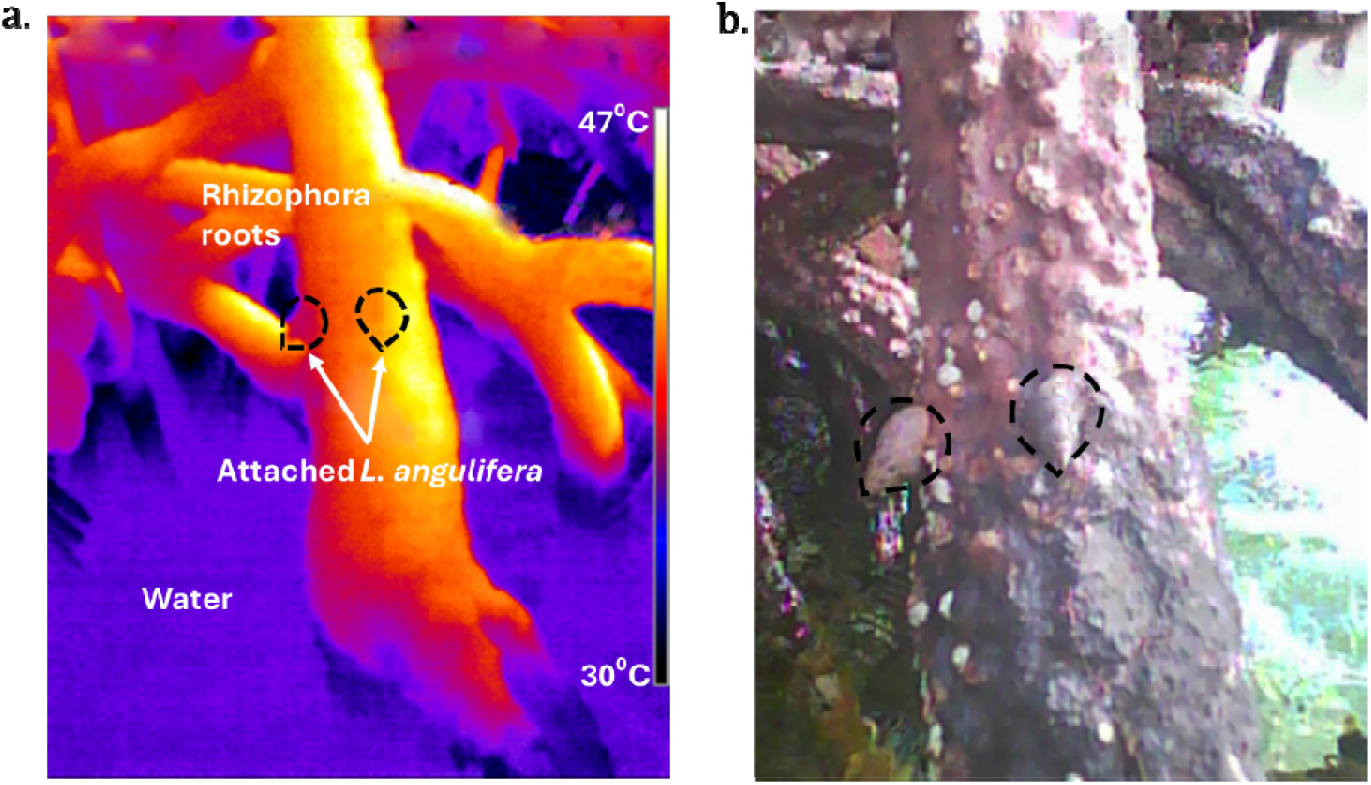
Panel a. Thermal image displaying the extreme thermal heterogeneity between the roots of *R. mangle* (red mangrove) and water below. Arrows point to two *L. angulifera* (circled) that glued above the waterline after feeding, with the snail on the left having a body temperature of 34°C, and the snail to the right, 40°C, checked by probing the internal body cavity with a thermal probe. Panel b. Normal photograph of the same snails for reference. Photographs taken on HTI Thermal Camera, (HTI instruments, China).

In this study we made three predictions: (1) *L. angulifera* microhabitat use is related to predation threat, with predation near the water’s surface being lower during the day, when *L. angulifera* are feeding, compared to at night. (2) *C. sapidus* has a higher warming tolerance, or difference between maximum habitat temperatures and upper thermal tolerance, than *L. angulifera*. (3) Sublethal heat exposure (near CT_max_) of L. angulifera reduces their capacity to behaviorally avoid predation and increase susceptibility to consumption. We tested these predictions using a combination of *in-situ* temperature monitoring, physiological and behavioral thermal trials, and a field out-planting experiment to compare the relative responses of interacting predator and prey to increasing habitat temperatures and extreme heat exposures.

## Materials and methods

### Study site and animal collection

*L. angulifera* and *C. sapidus* have an overlapping distribution from southern Florida to northern Brazil. Our study sites were in the red mangrove-dominated bays surrounding the Smithsonian Tropical Research Institute (STRI) on Colon Island, Bocas del Toro, Panama (9.351748N, -82.259182W). Collections were made between March – June 2024. All collections and experiments with *L. angulifera* and *C. sapidus* were approved by the Panama Ministry of Environment Department of Biodiversity under permit number ARB-062-2024. For all of the experiments below, we collected adult male (18-23 mm shell length) *L. angulifera* and male *C. sapidus* (70-90mm carapace width). Snails were collected by hand from a nearby mangrove patch, while crabs were collected using dip nets at 0.5-2 meter depth from a mangrove lagoon in the north part of the island (9°24’45.2”N 82°19’51.6”W) to avoid altering the predator abundance at the sites where we conducted the predation survey experiments.

Snails were maintained in small plastic reptile cages (18cm length x 35cm width x 22.5cm height) outdoors with ∼ 4cm depth of seawater on the bottom of the cages. Crabs were maintained in flow-through sea water aquariums at (29°C) (300L). Both species were kept for no longer than 48 hours.

### Environmental data

To understand the range of microhabitat temperatures that *L. angulifera* experiences in its natural habitat, we placed temperature loggers (iButtons, DS1921G, accuracy ±1°C, Maxim Integrated Products, USA) throughout mangrove roots and branches within the observed distribution of *L. angulifera* to record temperatures between April – June, 2024. iButtons were wrapped in Parafilm and then dipped in waterproof silicone, which closely matches the thermal properties of living snail tissue (Judge et al. 2018), and painted matte brown to reflect the color and texture of the exterior of *L. angulifera* shells (here after termed ‘biologgers’). Biologgers were retrieved every 2 – 3 weeks and the data were downloaded before resetting the biologgers. Live snails in the field were also periodically surveyed and body temperatures measured using a Sper Scientific temperature probe with a K-type thermocouple, to better understand the exact body temperatures snails experience compared with biologger recordings. To determine environmental temperatures for the crabs, water temperature was monitored at a depth of approximately 5 m (2 m off the bottom) on the instrument platform at the Bocas del Toro Research Station (BRS). Data were logged at 15 min intervals by data loggers attached to a Campbell Model 107 electronic temperature sensor. Because predation of *L. angulifera* occurs in shallower waters near the mangrove roots, we also set iButtons sealed in parafilm 0.25-0.5m below the water near the mangrove roots to determine if the microhabitat where predation occurred was warmer.

### Relating microhabitat use to predation vulnerability

To understand how sublethal heat stress could impact snail vulnerability to predators for *L. angulifera* on red mangroves, we conducted a restriction experiment to experimentally test how snail height on the mangroves is related to vulnerability to predation. Snails were restricted at 10 cm increments from the water line to a height of 50 centimeters on red mangrove prop roots of similar thickness and structure. Restriction was achieved by placing individual snails between two bands of copper tape wrapped around the mangrove prop root at each height, with the space in between band approximately 4cm, to allow snails horizontal but not vertical movement. Copper restriction is a common method used to restrict intertidal gastropods because snails have a strong aversion to copper and thus will not cross copper-covered surfaces (Gosselin & Chia 1995, Cubit 1984, Menge et al. 1993). After being placed between bands, snails were left in the field for a period of 24 hours. 10 snails were used per height for each trial (n =10 replicate snails per height + n = 8 trials, n = 80 snails per height total).

At the start of a survey, snails were set on the roots at dawn (6:00) and then individually checked before dusk (19:00) and then checked again the following morning (6:00). Snails found eaten at the 19:00 survey check were replaced with new snails to ensure the same number of snails present for both diurnal and nocturnal survey counts. The number of snails consumed before the 19:00 survey were counted as snails eaten during the day, and those consumed between 19:00 and 6:00 the following morning were considered eaten at night. Surveys occurred in a different bay than the one used for the sublethal heat exposure experiments to avoid altering predator behavior or preferences to certain roots. Seven separate trials were conducted from April – June 2024.

### Physiological thermal performance and upper thermal limit– resting heart rate

To compare *L. irrorata* (snail) and *C. sapidus* (crab) physiological thermal performance, we measured the resting heart rate (as a proxy for metabolic rate) in response to thermal ramping in the laboratory. We used a non-invasive photoplethysmography method (Burnett et al., 2013, Dwane et al. 2023), which uses an infrared sensor (CNY70, Vishay semiconductors, Germany) placed over the heart (dorsal carapace for crabs, or the body whorl of the snails’ snail) and attached using a thin layer of dental wax and super glue. After the sensor was attached, the animal was returned to an individual holding container for 24 hours to allow the glue to cure and the animal to rest before beginning the trials.

The infrared sensor was connected to an amplifier unit (AMP03, Newshift Lda., Portugal), which connects to a data acquisition system (USB-6001, National Instruments, USA) connected to a laptop running the NI-DAQExpress program (National Instruments, USA). Amplified signals were recorded at a sample frequency of 40 Hz. For trials using *L. angulifera*, snails were placed inside individual dry plastic bags (*n* = 3 snails per trial) and placed inside a water bath (VWR WB20 Digital Water Bath, VWR USA) set to 30°C. Snails were kept in air because *L. angulifera* is rarely found submerged and experiences its greatest thermal stress when emersed on mangrove roots. Bags were submerged except for their tops, which remained unsealed to allow gas exchange. To restrain the activity of the snails during the trials, a small piece of plastic mesh was attached to the shell that gently held the snail down to the bags surface, while it could still extend and retract its foot normally. Crabs are primarily subtidal; therefore, we measured crab heart rates while resting submerged in aerated water. Crabs were placed in glass beakers preheated to 28°C (*n* = 3 crabs per trial), which contained enough sea water to cover the crab completely, and the water was aerated using an air stone and bubbler. For restraining the crabs, thin wooden dowels were placed above the crab in the beakers and lightly pressed down until they held the crab in place.

Temperatures during the trials were monitored using a Sper Scientific 900005 thermal probe with a K-type thermocouple. For the snail trials, the probe was inserted inside an empty shell filled with aquarium silicone to mimic snail body temperatures and was placed in a separate bag alongside the snail trials. Because the snails were tested in air and did not immediately match air temperatures due to thermal inertia, this method can more accurately determine the body temperature of the snails than measurements of air or temperature alone. For the crab trials, the probe was placed directly into the water with the crab. Once equilibrium was reached, both crabs and snail trials underwent the same ramping rate of 1°C every 10 minutes (6 °C h^−^) until the animals reached cardiac flatline (cardiac lethal limit, or the upper thermal limit).

### Behavioral thermal performance and CT_max_

We measured behavioral thermal performance in *L. anguilifera* by assessing righting time-the time taken for a snail to right itself after being placed upside down - across a thermal ramp. Righting time is a common method used to asses behavioral performance across different taxa including marine and freshwater invertebrates (Lawrence & Cowell 1996 Brothers & McClintock 2015, Sherman 2015, DeWhatley & Alexander 2018, Collin et al. 2018), to estimate thermal optimum and CTmax (loss of righting responses, LRR) of behavioral activity. Snails were placed on individual glass dishes atop a warming plate (Monaco et al. 2016), starting from ∼30°C and ramping to 45°C. Righting time was tested for 12 snails at intervals of 2-3°C. To measure righting time, the heat plate was set to the starting temperature, and the snail was added once the surface temperature reached the target temperature. When the snail attached to the surface, it was left for 10 minutes to allow its body to equilibrate to the plate temperature, which was confirmed by probing a silicone-filled shell. After 10 minutes, the snail was gently lifted and flipped over. We then recorded the time it took for the snail to fully right itself (its foot reattached to the surface). After each trial, the plate was lightly misted to allow the snail to rehydrate and the heat plate was increased ∼2.5 degrees, allowing the snail to rest and equilibrate to the next trial temperature before repeating the trial again. The trials were repeated for individual snails at each temperature until the snails were incapable of righting themselves, which we recorded as the snails Loss of Righting Response (LRR) (McMahon 2001, Monaco et al. 2016). In some trials, snails did not emerge from the shells even at 30°C after 10 minutes, which we concluded as ‘shyness’ and these data were excluded from the analysis.

In a separate experiment, we measured CT_max_ in *L. angulifera* as the temperature at which snails lost their capacity to remain attached to a surface. Loss of attachment is an especially relevant metric for *L. angulifera* because the loss of muscular control can result in snails falling off the mangroves and into the water, where they can easily be consumed by aquatic predators. To measure CT_max_, snails were placed individually in a dry glass vial with a moistened paper towel to maintain humidity and allowed to attach to the vials before beginning (Marshall et al 2015). The vials were placed in a water bath set to the same ramping protocol as in the heart rate trials, 1℃ per 10 minutes (Monaco et al. 2016, Marshall & McQuad. 2011), roughly equivalent to the rate of warming snails can experience in their environment. CT_max_ for each snail was recorded as the temperature at which the snail lost control of its foot muscles and could no longer remain attached to the vial, verified by lightly tapping the vial to test whether the snail was attached. After the trials, snails were returned to their outdoor cages and monitored for 48 hours to confirm the animals survived the trial.

To assess CT_max_ in *C. sapidus*, we used the same ramping protocol as used for the heart rate trials for another set of 12 adult males. Crabs were added directly to the water bath set to 28ºC and monitored throughout the ramping of temperatures, until the crabs appeared to be unable to hold themselves upright or showed uncoordinated movements (loss of equilibrium). To confirm CT_max_ based on previously developed methods in portunid crabs, we turned over the crabs at these temperatures with prongs, and CT_max_ was determined if crabs were not able to right themselves (Vinagre et al 2015). Similar to the *L. angulifera* trials, water temperatures were immediately lowered after reaching CT_max_, and crab survival was monitored for 48 hours after trials to confirm survival. Data from crabs that perished during recovery were not included in the final analysis of CT_max_.

### The effects of sublethal heat exposure on predation vulnerability

To test the effects of sublethal heat stress on the susceptibility of *L. angulifera* to predation in the field, we conducted a field out-planting experiment, where snails were experimentally exposed to sublethal temperatures in the lab, before out-planting out into the field to monitor the impacts of heat exposure on susceptibility to predation in the mangroves. We chose sublethal exposure temperatures from 41°C - 45°C. They are well below the cardiac lethal limit (53.4°C), and at and below the CT_max_ (45.4°C SE). They also represent environmentally realistic temperatures of the extreme, 90^th^ percentile temperatures experienced in mangroves during the day for durations of up to 6 hours based on the biologger data (90^th^ percentile = 41.5°C, mean of 44.51°C, Fig. 3, Supplemental ST5).

For the exposure treatments, snails were collected 24 hours previous, marked with a unique ID (Beetag), and kept in plastic containers outside the lab in a shaded area to avoid acclimation to laboratory conditions. Snails were placed in glass vials with a moist paper towel to maintain humidity, and covered in parafilm with holes pierced at the top to maintain air flow into the vials. The vials were then added into a metal tray holder and placed into water baths set to 33°C, the approximate mean temperatures of biologgers in the field (mean = 33.97°C), with most of the vial being submerged to maintain an internal temperatures that reflected the surrounding water. Once the vials were added, the water bath temperature was raised from 33°C to its assigned treatment temperatures (41°C, 42°C, 43°C, 44°C, or 45°C) at an approximate rate of 1°C per 10 minutes and then held at the assigned temperature for six hours. Although this duration is longer than the duration of heat exposure at which we measured CT_max_ and cardiac lethal limit in our other experiments, preliminary trials confirmed that snails fully recovered from 6-hour exposure to the treatment temperatures. Target temperatures and rates of warming were confirmed by placing an empty snail shell filled with silicone in a vial in each treatment and checked using a Sper Scientific thermal prob with a K-type thermal couple. Temperatures were generally ±0.5°C from the target temperature.

Pre-exposures occurred at the same time each day (12:00 - 18:00), after which the snails were removed from the vials, added to plastic containers, and brought to the experimental site. Snails were then moved to a mangrove patch and placed on mangrove roots chosen at random among 20 marked roots that were identified previously to be similar in size, structure, and epiphyte growth to minimize any difference that could lead to difference in predator access to snails. For each trial, snails were placed ∼ 30 minutes before dusk at the same heights on the roots (at the water line) and allowed to attach. To prevent snail escape, copper tape was wrapped around the trunk of the mangrove 1 meter above the roots, so snails could move outside of the range of subtidal predation (Fig 2) but not leave the experimental area.

**Fig. 2.**
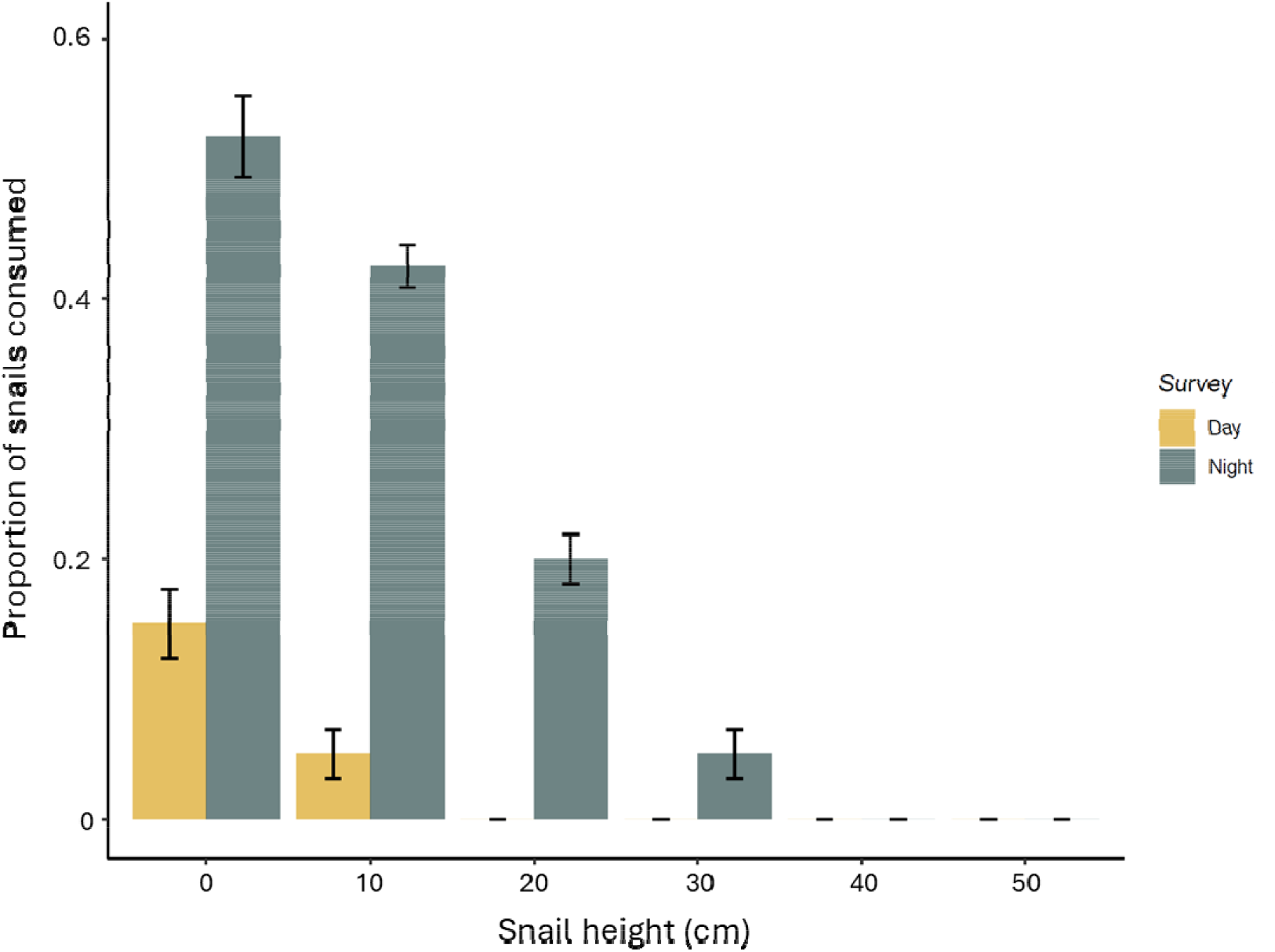
The proportion of *Littoraria angulifera* consumed at each height above the water line during the day (gold) and night (grey).

Snails were then checked the following morning after dawn, and the roots searched for all remaining snails. Each trial had n=20 snails per treatment temperature. Snails were placed in groups of 5 on a root. Each temperature was tested 3 times with 20 snails in each trial, totaling n=60 snails for each temperature treatment. Because predation events could not be confirmed visually by this method, we determined a ‘probability of presumed predation’ based on the number of snails missing from the roots in the morning survey.

### Statistical analysis

For the snail restriction experiments, we constructed a generalized linear mixed model (GLMM) to test if predation is highest at night compared to during the day, and the relative vulnerability of predation based on heights. To estimate the effects of snail height, time of survey, and their interaction (fixed effects), on the probability of mortality (response variable), with survey number included in the model as a random effect to account for non-independence. We used a binomial distribution because the probability of mortality is proportion data. We used the “glmer” function from the *lme4* package. Model assumptions were checked and met using the *DHARMA* package. We then ran a post-hoc pairwise comparison between height and time with a Tukey HSD test using the *emmeans* package.

For testing the impacts of acute heat stress on snail vulnerability to predation, we again used a GLMM, with temperature treatment as a fixed effect, and trial number, root, and group that the snails were assigned treated as random effects.

Cardiac and righting time thermal performance curves (TPCs) were analyzed using the R package *rTPC* following the pipeline by Padfield et al. (2021). We wanted to test how temperature of peak activity (T_peak_, commonly referred to in the literature as T_opt_ or thermal optimum) and upper thermal limits (cardiac lethal limit) differed between crabs and snails, which we then extracted to investigate how close each species was living to these parameters in its habitat (*warming tolerance* = *90*^*th*^ *percentile temperatures* – *upper thermal limit*). We also wanted to extract other parameters that are useful for predicting vulnerability to warming, such as thermal optimum and thermal safety margin (the difference between thermal optimum and upper thermal limit. This has also been referred to as the difference between maximum habitat temperatures and thermal optimum (Deutch et al. 2008). For each species, we thus combined individual TPCs (n=12 for each species) to create a ‘species-level TPC’. We then chose 9 models from the package that could provide suitable fits to our TPCs based on their previous use on similar taxa and traits (see table ST6 – ST8 for model types and parameters). To test for model fit, we used non-linear least squares regression, and fit was determined by choosing the models with the lowest Akaike Information Criterion corrected for small sample size (AICc) across the models (Supplemental tables ST6 – ST8), and confirmed by reviewing the visual fit of the curves. We used the same model, Sharpsschoolfield, for heart rate TPCS between *L. angulifera* and *C. sapidus* because it had the lowest AICc for *C. sapidus* and second lowest for *L. angulifera*, and best visual fit for both, making parameters comparable between species. We then used weighted residual bootstrapping to assess model uncertainty (Pallarés et al. 2021, Padfield et al. 2021). TPC parameters for each TPC were then generated with a 95% confidence interval. We tested for differences in CT_max_ between *C. sapidus* and *L. angulifera* using a Student’s t-test, which assumes normality and homogeneity of data between the two data sets, which was confirmed using a Shapiro-Wilk test and an F-test, respectively. All data analyses were performed using R version 4.5.2 (R Core Team, 2025).

## Results

### Relating prey height to vulnerability to predation between day and night

The proportion of snails consumed differed significantly between survey times (*χ*^*2*^ = 44.25, *df* = 1, *p* < 0.001) and among heights (*χ*^*2*^ = 39.23, *df* = 5, *p* < 0.001). Presumed predation was significantly higher at night between 0 – 30 cm on the roots compared to daytime surveys, and predation was highest at the water line and gradually reduced where snails experienced no predation at or above 20cm during the day, and at or above 40 cm at night (Fig. 2, ST2).

### Cardiac thermal performance relative to habitat temperatures

*L. angulifera* had a mean upper thermal limit (cardiac lethal limit) of 53.4°C, and thermal optimum of 45.9°C. The average temperatures of the 90^th^ percentile temperatures from biologgers were 44.5°C. (ST5), indicating a warming tolerance of approximately 8.9°C (Table 1; Fig. 3a). *C. sapidus* have a cardiac lethal limit of 44.5°C, and a thermal optimum of 40.9°C (Table 1; Fig. 3b). The average temperatures of the 90^th^ percentile temperatures from water temperatures below the mangroves was 32.7°C, indicating a warming tolerance of approximately degrees of 11.8°C (ST5).

**Table 1.**
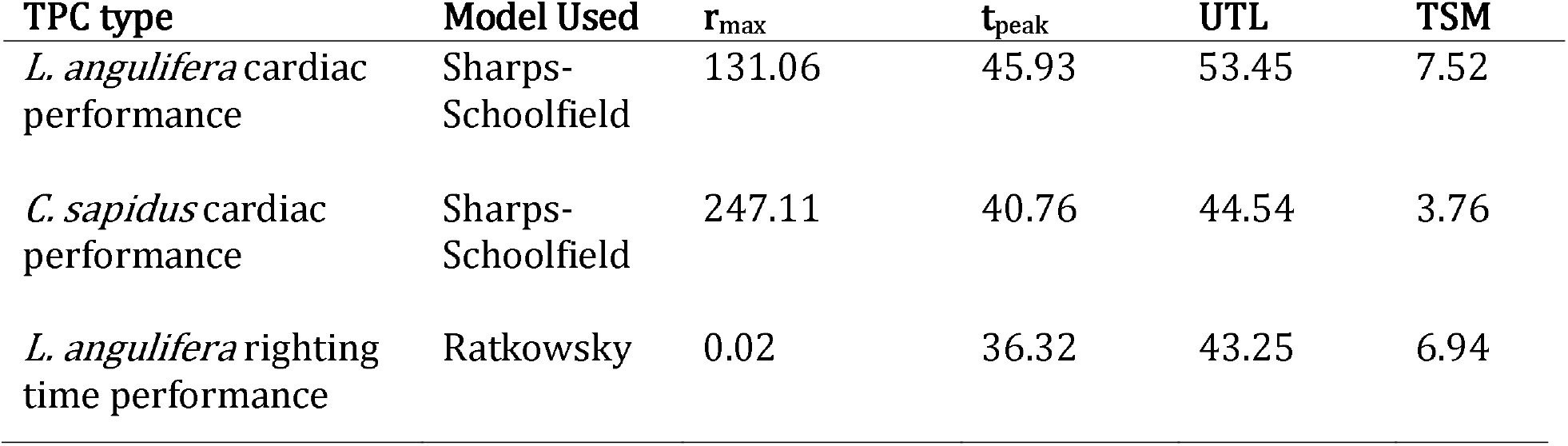
Parameter estimates from the cardiac and behavior thermal performance curves. r_max_: maximum heart rate; T_peak_: temperature of peak performance (°C); UTL (upper thermal limit = cardiac lethal limit); upper thermal limit of performance; T_br_: thermal breadth (°C); TSM (thermal safety margin): Difference (°C) between T_peak_ and UTL.

**Fig. 3.**
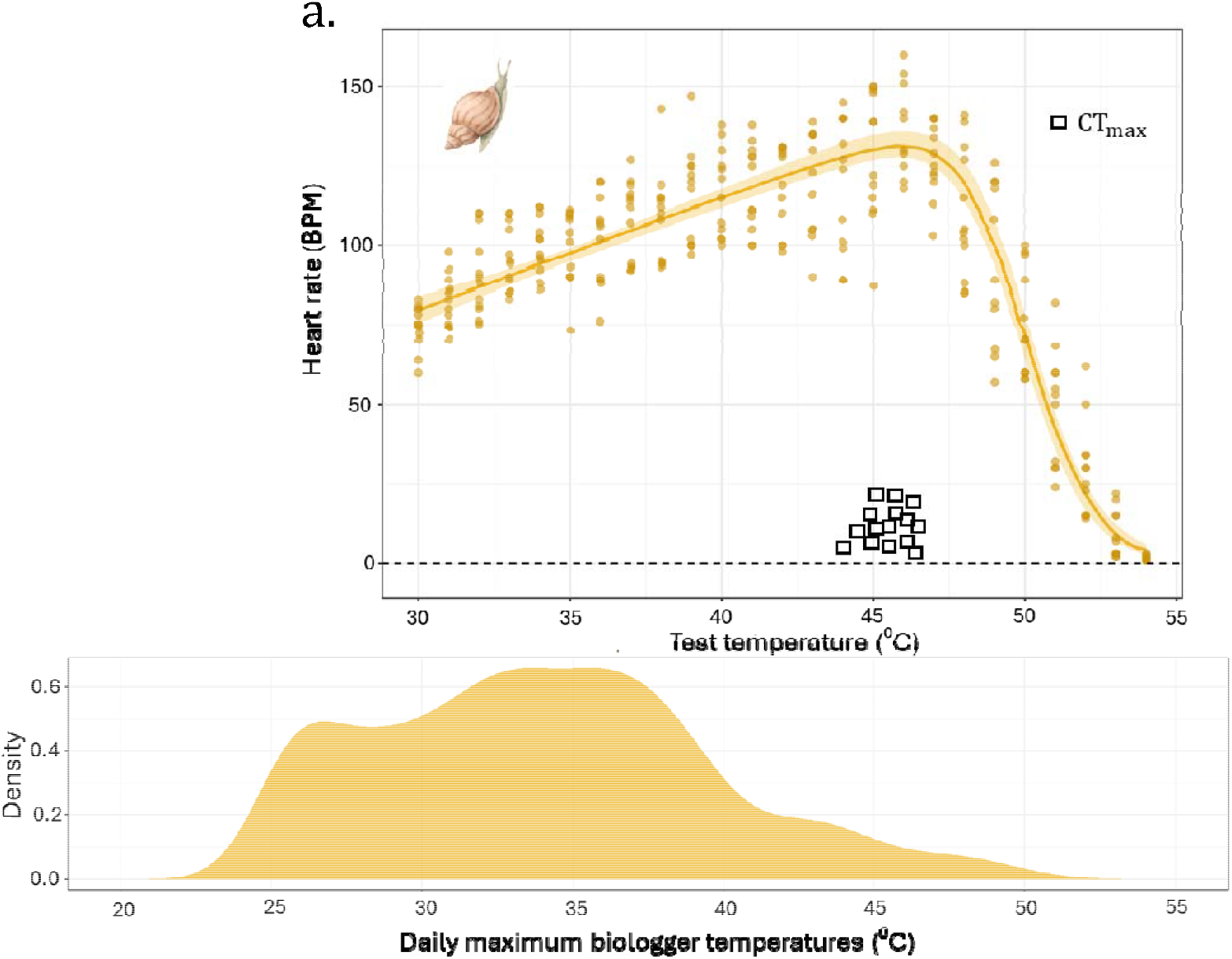
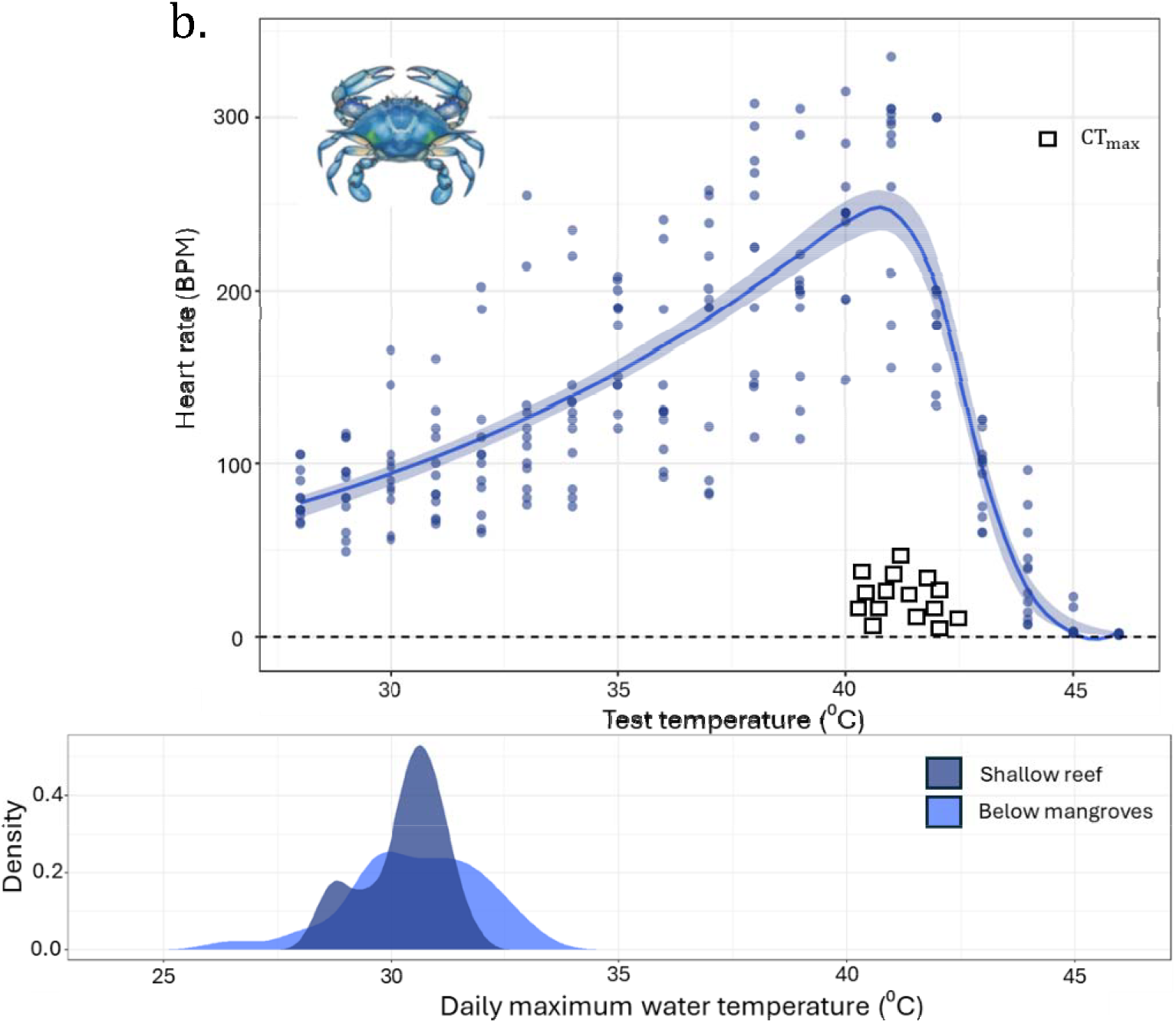
The cardiac thermal performance of *Littoraria angulifera* (panel a, gold) and *Callinectes sapidus* (panel b, blue) and daily maximum temperatures within each species’ respective habitats. Squares represent each species CT_max_ (loss of attachment in snails; loss of equilibrium in crabs; Supplemental S1, ST9). Points are jittered for easier visualization and comparison to the cardiac lethal limit (upper thermal limit). Density distributions below the thermal performance curves are the daily maximum temperatures recorded during the study period from March – June 2025. The line represents model output for each species performance across temperatures, and shaded areas represent 95% confidence interval for the fitted TPC models.

### Critical thermal maximum and righting time thermal performance

Critical thermal maximum (CT_max_) was lower than the cardiac lethal limit for both species. *L. angulifera* exhibited the greatest difference, where mean CT_max_ for *L. angulifera* was 45.4 ± 0.51°C (mean ± SD), 8°C lower than the cardiac lethal limit (upper thermal limit) (Fig. 3a, Supplementary Figure S1, ST9). *C. sapidus* had a mean CT_max_ of 41.46 ± 0.66°C, 3.04°C lower than upper thermal limit (Fig. 3b, Supplementary Figure S1, ST9). For righting timing thermal performance of *L. angulifera*, the thermal optimum was 36.3°C, with a loss of righting response at 43.25°C (Fig. 4; Table 1).

**Fig. 4.**
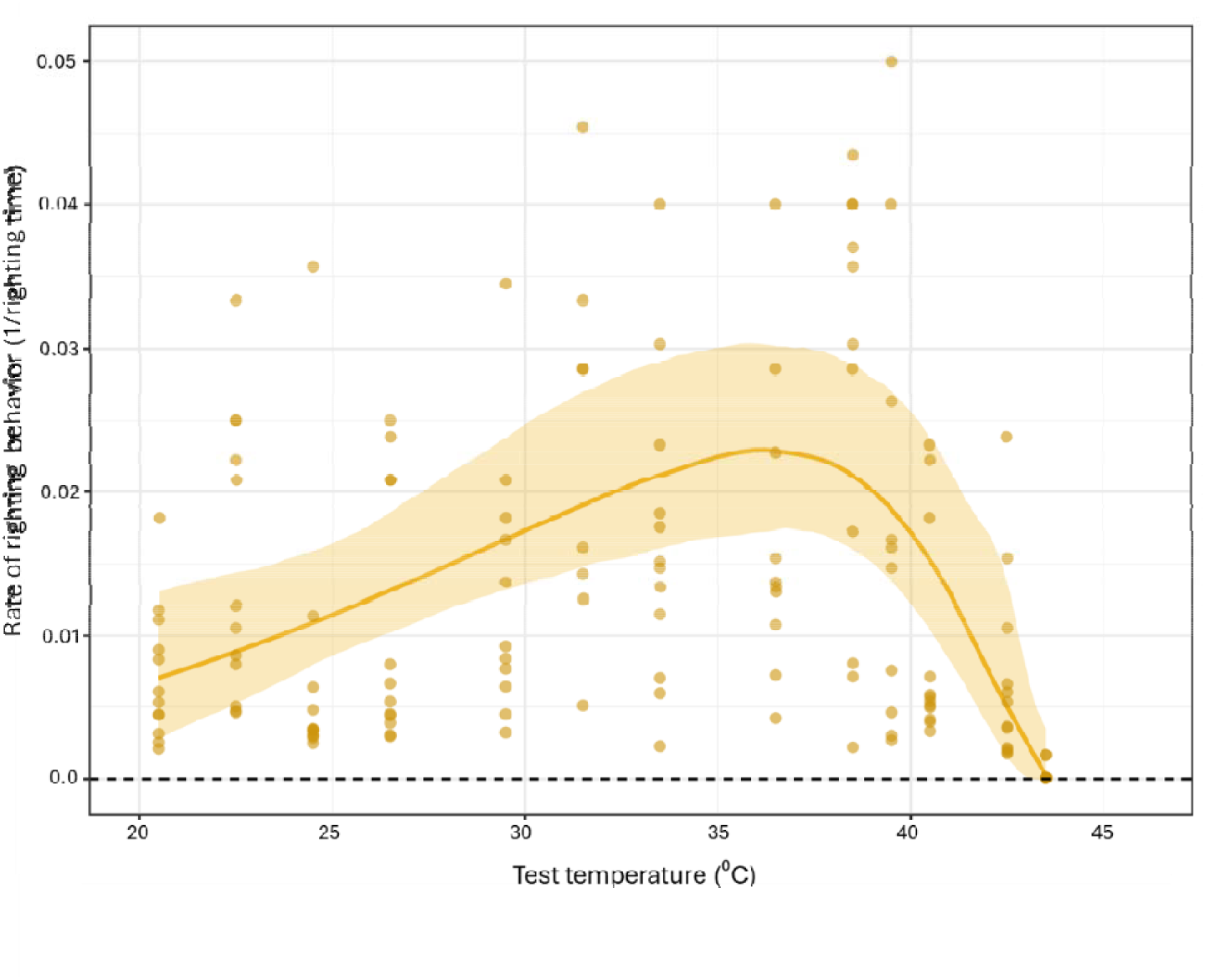
Thermal performance curve of righting time performance of *L. angulifera*. Extreme outliers (5% of data) were cropped from the graph to improve the visibility of the shape of the curve, but all data was included for generating the models. The line represents the model output for performance across temperatures, and shaded areas represent 95% confidence interval for the fitted TPC model.

**Fig. 5.**
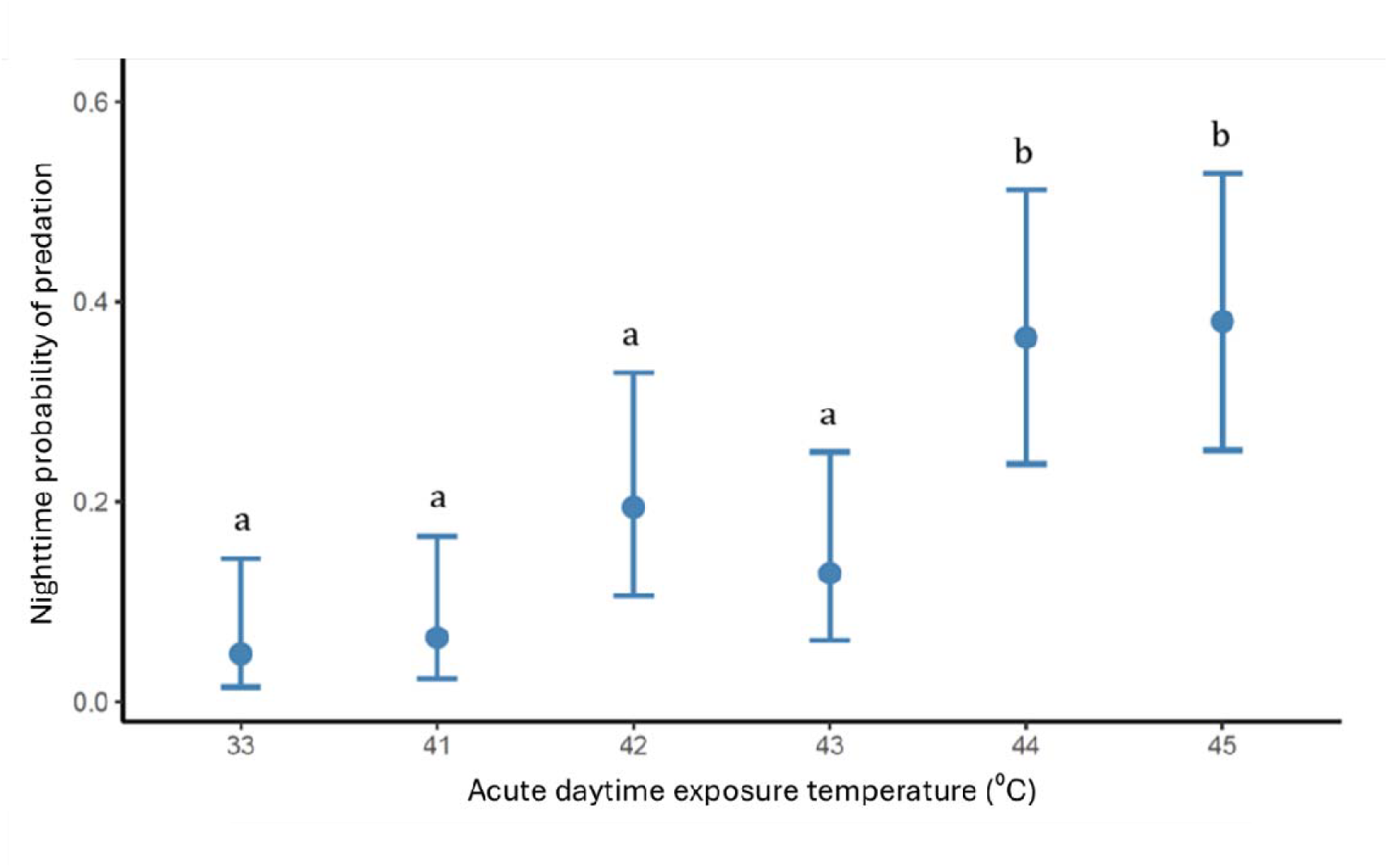
Impacts of acute heat stress exposure on the proportion of *L. angulifera* missing in the out-planting experiment. Different letters represent significant difference in presumed predation among treatments based on post-hoc tests.

### Impacts of acute exposure on snail vulnerability to predation

The proportion of snails that were predated differed strongly between temperature treatments (*χ*^*2*^ = 33.07, *df* = 5, *p* < 0.001). Snails that were exposed to temperatures of 44°C and 45°C experienced higher proportion of predation (36 ± 7% & 38 ± 7% respectively; mean ± SE) compared to control (33°C, 5 ± 2%) treatments. Although predation was higher in lower heat exposure treatments than control (41°C, 6.4 ± 3%, 42°C, 19 ± 6%, and 43°C 13 ± 5%), they were not significantly different than proportion of snails consumed at control temperatures (Supplemental ST3 and ST4).

## Discussion

Ecological communities are increasingly exposed to extreme temperatures that are threatening species persistence and altering ecological interactions (Urban 2024), which may in part be impacted by asymmetries in thermal sensitivity between interacting species and resulting effects of these asymmetries across trophic levels (Zarnetske et al. 2012). In our study system, we found that while both predator (*C. sapidus*) and prey (*L. angulifera*) exhibit high upper thermal limits, *C. sapidus* are less at risk of encountering acute thermal stress in the field due to its aquatic habitat acting as a refugium against thermal extremes.

In contrast, we found that *L. anguilifera*, despite possessing higher lethal limits, is more likely to encounter field temperatures close to sublethal thresholds that impair behavioral and physiological performance. Furthermore, by experimentally exposing prey to heat stress and out-planting them into the field, we found that exposure to temperatures near these sublethal thresholds significantly increased predation risk by compromising preys’ ability to behaviorally avoid predators during the recovery window. This provides striking evidence that sublethal assays of physiological and behavioral stress can correspond to severe ecological costs in nature, even in the absence of direct mortality. Our study also highlights that we may mischaracterize species vulnerability to warming if we assume interacting species are experiencing the same levels of thermal stress.

Predator-prey interactions that occur at the interface between aquatic and terrestrial habitats are ubiquitous across taxa and ecosystems (Dill et al. 1977, Shoop & Ruckdeschel 1990. Duncan & Szelistowski 1998, Amr et al. 2018, Ohba 2019, Jawad et al. 2024), and the strength of these interactions can regulate ecological communities (Silliman & Bertness 2001, Matassa et al. 2017). Predators that are primarily subaquatic should experience less variation and extremes in temperatures compared to their emersed prey, creating a mismatch in thermal habitats. This can have important consequences for predicting the vulnerability and resilience of these ecosystems to climate warming, and for predator-prey interactions under global change more generally. Previous work in temperate rocky shores has suggested that subaquatic predators (sea stars) are living below their thermal optimum, while their emersed prey (mussels) are already living close to physiologically stressful conditions (Monaco et al. 2016). Such variation in exposure to thermal extremes and thermal performance between subaquatic predators and prey may provide a mechanistic explanation for observed localized extirpations of prey, but not of their main predators, as these habitats warm (Harley et al. 2010). Our study suggests that mobile prey in tropical intertidal ecosystems may be similarly faced with increased vulnerability to warming relative to their predators, but the mechanism may be driven by the unique decoupling of behavioral and physiological tolerance to heat stress in the prey. Although snails have a high physiological tolerance to heat stress, susceptibility to predation was high for prey that had previous exposure to temperatures a full 10°C below lethal limits. In the tropics, mangroves are considered more thermally benign habitats compared to rocky shores (Marshall et al. 2015). The high physiological tolerance, but significantly lower behavioral tolerance (CT_max_) of littorines may reflect adaptation to rocky shores before colonizing mangroves (Marshall et al. 2015), where snails in rocky shores would need to ‘sit and physiologically fight’ extreme temperatures, but not necessarily be active during or after exposures to stressful temperatures. In mangroves, however, snails must climb a substantial distance above the water line after daytime heat has dissipated to avoid nighttime predation (Fig. 2). Higher susceptibility to predation after acute heat exposure may thus be due to hampered activity or immobilization during recovery, inhibiting movement up the mangroves to avoid subtidal predation at night. Our results emphasize the importance of incorporating local ecological contexts, such as when and where prey are most vulnerable to predation, for understanding how increasing heat stress may impact species vulnerability, both physiologically and as a result of shifting species interactions.

Our field surveys suggest that snails are active near the water line during the day but climb mangroves at dusk to avoid nocturnal predation which can be high near the water at night. However, our study did not test alternative explanations for the snails’ use of lower microhabitats during the day, such as the benefits of foraging when temperatures are closer to thermal optimum for activity. Indeed, the behavioral TPC for the snails suggests that the T_peak_ of activity is approximately 36.3°C, temperatures that are only experienced during the day and on warmer portions of mangrove roots (Fig. 2). Being active in such conditions may increase snails’ grazing rate but come at the risk of increased exposure to acute stress. Previous work has suggested that temperate intertidal snails may prefer cooler microhabitats well below their thermal optimums to avoid risk of heat stress exposure later in the day (Tepler et al. 2011), yet no studies to our knowledge have investigated thermal preference in relation to feeding and growth rate in littorines (but see Atkins et al. 2022). Such tradeoffs may be common in intertidal species, where the access to food may trade-off with thermoregulation (Hayford et al. 2018).

Heat waves are increasing in frequency and intensity across both terrestrial and marine ecosystems, and recent work has suggested terrestrial and marine heat waves may be increasingly linked (Pathmeswaran et al. 2022). Because the water temperatures of shallow water habitats will be a function of both sea surface and air temperatures, it is difficult to extrapolate how species that interact between aquatic and air environments will be differentially exposed to climate events. Water temperatures immediately below the mangrove roots were only a few degrees warmer than the deeper reef, and still well below the thermal tolerance and T_peak_ of the crabs cardiac activity. Indeed, our study supports previous findings that portunid (swimming) crabs may have high thermal tolerance and performance relative to their habitat temperatures, predicting persistence and habitat expansion of the species as the climate warms (Marchessaux et al. 2024). However, aquatic predators may still be increasingly vulnerable to warming events relative to emersed prey because warming water, especially in calm estuarine environments, has reduced oxygen availability, which may limit activity and heat tolerance in aquatic species (Pörtner et al. 2010, Coates et al. 2022). Indeed, punctuated hypoxia events can be lethal for aquatic invertebrates alongside heat waves, and the two can co-vary in the Bocas Archipelago (Lucey et al. 2023).

Because we only used one predator species in our study, we do not know if other predators of *L. angulifera* have similar heat tolerance. Molluscivorous fish that are known predators of littorines, such as puffer fish (family Tetraodontidae), have substantially lower heat tolerance than crabs, based on previously published data on thermal tolerances (Shultz et al. 2016). The high tolerance of the crabs may reflect their life history, where they can occupy a suite of shallow water habitats across their life stages and activity (Johnson 2022), where exposures to high temperatures and periods of emersion or becoming stranded in low tidal habitats could still impose selection for high physiological thermal tolerance. Integrating observations of species habitat use and thermal exposures across a wider portion of species life stages will be essential for more comprehensive estimation of species responses to climate warming. In addition, comparing thermal traits among multiple predator and prey within the same habitat, including a wider taxonomic diversity with different life histories, can provide insight into the contexts and traits that may predict which types of interactions may be most vulnerable or resilient to global change, which will have important implications for subsequent impacts on the community as a whole (Kroeker & Sanford 2022, Urban et al. 2017).

Integrating physiological and behavioral thermal sensitivity is a useful framework for predicting the responses of interacting species to climate warming. Including such data alongside the ecological context of the interactions, such as when heat stress and the interaction is strongest, will add important nuance. Nocturnal predator and/or prey activity is common across ecosystems, yet little is known about how increasing daytime temperatures can impact interactions at night. We show that CT_max_ can be a useful metric to predict when prey can become ecologically vulnerable to predation, and that the impacts can carry over into nocturnal activities, emphasizing that the effects of sublethal stress may be a severally underappreciated consequence of climate warming. Sublethal effects could be especially important for susceptibility to predation in ecosystems where prey are differentially exposed to thermally stressful conditions compared to their predators. Taken together, our work demonstrates that both nuanced physiological data, and an understanding of ecological context will be critical to anticipating biological responses to global change.

## Acknowledgments

We would like to thank the staff at the Smithsonian Tropical Research Institute, Bocas del Toro Research Station, for extensive help with logistics and support for this study. We thank Ana Salgado, Kyle Harms, and Michael Hellberg for helpful suggestions in preparing the manuscript and analysis. Equipment and stay at the station was funded by the STRI Short-Term Fellowship Award, awarded to W.A.J. M.W.K was supported by NSF-IOS 2154283. C.D was supported by NSF-OCE-2023571 while at the University of Massachusetts Amherst.

## Author contributions

All authors conceived the ideas and designed methodology; W.A.J collected the data; W.A.J and C.D analyzed the data; W.A.J led the writing of the manuscript. All authors contributed to drafts of the manuscript and final approval for publication.

## Data availability statement

All data and scripts will be placed in a public repository (Zenodo) and made available upon acceptance.

