## Supplemental material for "Daytime heat exposure increases nighttime predation risk in a mangrove gastropod"

ST1 Summary of GLM for predation height restriction surveys (put number of observations/trials here in caption)

| *Random effects* |  |  |  |  |
| --- | --- | --- | --- | --- |
| **Groups** | **Name** | **Variance** | **Std.Dev.** |  |
| Trial | (Intercept) | 0 | 0 |  |
| Number of observations: 96, groups: trial 8 |  |  |  |  |
| *Fixed effects* |  |  |  |  |
| **Term** | **Estimate** | **Standard error** | **Z-value** | **P-value** |
| (Intercept) | -1.9459 | 0.3381 | -5.756 | **8.61E-09** |
| SurveyNight | 1.9459 | 0.4053 | 4.801 | **1.58E-06** |
| Snail.verticle.height10 | -0.9985 | 0.6144 | -1.625 | 0.104 |
| Snail.verticle.height20 | -22.1556 | 19144.23 | -0.001 | 0.999 |
| Snail.verticle.height30 | -22.1556 | 19144.02 | -0.001 | 0.999 |
| Snail.verticle.height40 | -22.1556 | 19144.05 | -0.001 | 0.999 |
| Snail.verticle.height50 | -22.1556 | 19144.05 | -0.001 | 0.999 |
| SurveyNight:Snail.verticle.height10 | 0.6962 | 0.6918 | 1.006 | 0.314 |
| SurveyNight:Snail.verticle.height20 | 20.7693 | 19144.23 | 0.001 | 0.999 |
| SurveyNight:Snail.verticle.height30 | 19.4476 | 19144.02 | 0.001 | 0.999 |
| SurveyNight:Snail.verticle.height40 | -1.9459 | 27073.78 | 0 | 1 |
| SurveyNight:Snail.verticle.height50 | -1.9459 | 27073.78 | 0 | 1 |

ST2 Post hoc table

| **Contrast** | **Estimate** | **SE** | **df** | **z.ratio** | **p.value** |
| --- | --- | --- | --- | --- | --- |
| Day Snail.verticle.height0 - Night Snail.verticle.height0 | -1.94591 | 0.405322 | Inf | -4.8009 | **0.0001** |
| Day Snail.verticle.height0 - Day Snail.verticle.height10 | 0.998529 | 0.614364 | Inf | 1.625304 | 0.900144 |
| Day Snail.verticle.height0 - Night Snail.verticle.height10 | -1.64363 | 0.406739 | Inf | -4.04099 | **0.0031** |
| Day Snail.verticle.height0 - Day Snail.verticle.height20 | 22.15563 | 19144.23 | Inf | 0.001157 | 1 |
| Day Snail.verticle.height0 - Night Snail.verticle.height20 | -0.55962 | 0.438646 | Inf | -1.27578 | 0.982155 |
| Day Snail.verticle.height0 - Day Snail.verticle.height30 | 22.15561 | 19144.02 | Inf | 0.001157 | 1 |
| Day Snail.verticle.height0 - Night Snail.verticle.height30 | 0.76214 | 0.57238 | Inf | 1.331528 | 0.975142 |
| Day Snail.verticle.height0 - Day Snail.verticle.height40 | 22.15561 | 19144.05 | Inf | 0.001157 | 1 |
| Day Snail.verticle.height0 - Night Snail.verticle.height40 | 22.15561 | 19144.05 | Inf | 0.001157 | 1 |
| Day Snail.verticle.height0 - Day Snail.verticle.height50 | 22.15561 | 19144.05 | Inf | 0.001157 | 1 |
| Day Snail.verticle.height0 - Night Snail.verticle.height50 | 22.15561 | 19144.05 | Inf | 0.001157 | 1 |
| Night Snail.verticle.height0 - Day Snail.verticle.height10 | 2.944439 | 0.559605 | Inf | 5.261637 | **9.27E-06** |
| Night Snail.verticle.height0 - Night Snail.verticle.height10 | 0.302281 | 0.318042 | Inf | 0.950442 | 0.998571 |
| Night Snail.verticle.height0 - Day Snail.verticle.height20 | 24.10154 | 19144.23 | Inf | 0.001259 | 1 |
| Night Snail.verticle.height0 - Night Snail.verticle.height20 | 1.386294 | 0.357946 | Inf | 3.87292 | 0.006054 |
| Night Snail.verticle.height0 - Day Snail.verticle.height30 | 24.10152 | 19144.02 | Inf | 0.001259 | 1 |
| Night Snail.verticle.height0 - Night Snail.verticle.height30 | 2.70805 | 0.51316 | Inf | 5.277204 | **8.52E-06** |
| Night Snail.verticle.height0 - Day Snail.verticle.height40 | 24.10152 | 19144.05 | Inf | 0.001259 | 1 |
| Night Snail.verticle.height0 - Night Snail.verticle.height40 | 24.10152 | 19144.05 | Inf | 0.001259 | 1 |
| Night Snail.verticle.height0 - Day Snail.verticle.height50 | 24.10152 | 19144.05 | Inf | 0.001259 | 1 |
| Night Snail.verticle.height0 - Night Snail.verticle.height50 | 24.10152 | 19144.05 | Inf | 0.001259 | 1 |
| Day Snail.verticle.height10 - Night Snail.verticle.height10 | -2.64216 | 0.560632 | Inf | -4.71282 | 0.000154 |
| Day Snail.verticle.height10 - Day Snail.verticle.height20 | 21.1571 | 19144.23 | Inf | 0.001105 | 1 |
| Day Snail.verticle.height10 - Night Snail.verticle.height20 | -1.55814 | 0.584194 | Inf | -2.66717 | 0.242741 |
| Day Snail.verticle.height10 - Day Snail.verticle.height30 | 21.15708 | 19144.02 | Inf | 0.001105 | 1 |
| Day Snail.verticle.height10 - Night Snail.verticle.height30 | -0.23639 | 0.690283 | Inf | -0.34245 | 1 |
| Day Snail.verticle.height10 - Day Snail.verticle.height40 | 21.15708 | 19144.05 | Inf | 0.001105 | 1 |
| Day Snail.verticle.height10 - Night Snail.verticle.height40 | 21.15708 | 19144.05 | Inf | 0.001105 | 1 |
| Day Snail.verticle.height10 - Day Snail.verticle.height50 | 21.15708 | 19144.05 | Inf | 0.001105 | 1 |
| Day Snail.verticle.height10 - Night Snail.verticle.height50 | 21.15708 | 19144.05 | Inf | 0.001105 | 1 |
| Night Snail.verticle.height10 - Day Snail.verticle.height20 | 23.79926 | 19144.23 | Inf | 0.001243 | 1 |
| Night Snail.verticle.height10 - Night Snail.verticle.height20 | 1.084013 | 0.35955 | Inf | 3.014921 | 0.1041 |
| Night Snail.verticle.height10 - Day Snail.verticle.height30 | 23.79924 | 19144.02 | Inf | 0.001243 | 1 |
| Night Snail.verticle.height10 - Night Snail.verticle.height30 | 2.405769 | 0.51428 | Inf | 4.677935 | **0.000183** |
| Night Snail.verticle.height10 - Day Snail.verticle.height40 | 23.79924 | 19144.05 | Inf | 0.001243 | 1 |
| Night Snail.verticle.height10 - Night Snail.verticle.height40 | 23.79924 | 19144.05 | Inf | 0.001243 | 1 |
| Night Snail.verticle.height10 - Day Snail.verticle.height50 | 23.79924 | 19144.05 | Inf | 0.001243 | 1 |
| Night Snail.verticle.height10 - Night Snail.verticle.height50 | 23.79924 | 19144.05 | Inf | 0.001243 | 1 |
| Day Snail.verticle.height20 - Night Snail.verticle.height20 | -22.7152 | 19144.23 | Inf | -0.00119 | 1 |
| Day Snail.verticle.height20 - Day Snail.verticle.height30 | -2.32E-05 | 27073.88 | Inf | -8.57E-10 | 1 |
| Day Snail.verticle.height20 - Night Snail.verticle.height30 | -21.3935 | 19144.23 | Inf | -0.00112 | 1 |
| Day Snail.verticle.height20 - Day Snail.verticle.height40 | -1.89E-05 | 27073.91 | Inf | -6.99E-10 | 1 |
| Day Snail.verticle.height20 - Night Snail.verticle.height40 | -1.89E-05 | 27073.91 | Inf | -6.99E-10 | 1 |
| Day Snail.verticle.height20 - Day Snail.verticle.height50 | -1.89E-05 | 27073.91 | Inf | -6.99E-10 | 1 |
| Day Snail.verticle.height20 - Night Snail.verticle.height50 | -1.89E-05 | 27073.91 | Inf | -6.99E-10 | 1 |
| Night Snail.verticle.height20 - Day Snail.verticle.height30 | 22.71522 | 19144.02 | Inf | 0.001187 | 1 |
| Night Snail.verticle.height20 - Night Snail.verticle.height30 | 1.321756 | 0.539869 | Inf | 2.448291 | 0.37324 |
| Night Snail.verticle.height20 - Day Snail.verticle.height40 | 22.71523 | 19144.05 | Inf | 0.001187 | 1 |
| Night Snail.verticle.height20 - Night Snail.verticle.height40 | 22.71523 | 19144.05 | Inf | 0.001187 | 1 |
| Night Snail.verticle.height20 - Day Snail.verticle.height50 | 22.71523 | 19144.05 | Inf | 0.001187 | 1 |
| Night Snail.verticle.height20 - Night Snail.verticle.height50 | 22.71523 | 19144.05 | Inf | 0.001187 | 1 |
| Day Snail.verticle.height30 - Night Snail.verticle.height30 | -21.3935 | 19144.02 | Inf | -0.00112 | 1 |
| Day Snail.verticle.height30 - Day Snail.verticle.height40 | 4.26E-06 | 27073.75 | Inf | 1.58E-10 | 1 |
| Day Snail.verticle.height30 - Night Snail.verticle.height40 | 4.26E-06 | 27073.75 | Inf | 1.58E-10 | 1 |
| Day Snail.verticle.height30 - Day Snail.verticle.height50 | 4.26E-06 | 27073.75 | Inf | 1.58E-10 | 1 |
| Day Snail.verticle.height30 - Night Snail.verticle.height50 | 4.26E-06 | 27073.75 | Inf | 1.58E-10 | 1 |
| Night Snail.verticle.height30 - Day Snail.verticle.height40 | 21.39347 | 19144.05 | Inf | 0.001118 | 1 |
| Night Snail.verticle.height30 - Night Snail.verticle.height40 | 21.39347 | 19144.05 | Inf | 0.001118 | 1 |
| Night Snail.verticle.height30 - Day Snail.verticle.height50 | 21.39347 | 19144.05 | Inf | 0.001118 | 1 |
| Night Snail.verticle.height30 - Night Snail.verticle.height50 | 21.39347 | 19144.05 | Inf | 0.001118 | 1 |
| Day Snail.verticle.height40 - Night Snail.verticle.height40 | -1.11E-15 | 27073.78 | Inf | -4.10E-20 | 1 |
| Day Snail.verticle.height40 - Day Snail.verticle.height50 | 6.39E-14 | 27073.78 | Inf | 2.36E-18 | 1 |
| Day Snail.verticle.height40 - Night Snail.verticle.height50 | -2.00E-14 | 27073.78 | Inf | -7.38E-19 | 1 |
| Night Snail.verticle.height40 - Day Snail.verticle.height50 | 6.51E-14 | 27073.78 | Inf | 2.40E-18 | 1 |
| Night Snail.verticle.height40 - Night Snail.verticle.height50 | -1.89E-14 | 27073.78 | Inf | -6.97E-19 | 1 |
| Day Snail.verticle.height50 - Night Snail.verticle.height50 | -8.39E-14 | 27073.78 | Inf | -3.10E-18 | 1 |

ST3 Summary of GLM for heat stress out planting surveys

| *Random effects* | |  |  |  |
| --- | --- | --- | --- | --- |
| Groups . | Name | Variance | Std. Dev. |  |
| Root | (Intercept) | 2.16E-02 | 1.47E-01 |  |
| Group | (Intercept) | 8.03E-02 | 2.83E-01 |  |
| Trial | (Intercept) | 6.66E-10 | 2.58E-05 |  |
| Number of obs: 72, groups: Root, 24; Group, 4; Trial, 3 | | | | |
| *Fixed effects* | |  |  |  |
| **Term** | **Estimate** | **Standard Error** | **Z-value** | **P-value** |
| (Intercept) | -2.9901 | 0.6176 | -4.842 | **1.29E-06** |
| Temperature41 | 0.3111 | 0.7971 | 0.39 | 0.696358 |
| Temperature42 | 1.5698 | 0.6841 | 2.295 | **0.021743** |
| Temperature43 | 1.0768 | 0.7134 | 1.509 | 0.131197 |
| Temperature44 | 2.4335 | 0.6599 | 3.688 | **0.000226** |
| Temperature45 | 2.5026 | 0.658 | 3.803 | **0.000143** |

ST4 Summary of post hoc comparison

| **Contrast** | **Odds ratio** | **SE** | **df** | **null** | **z.ratio** | **p.value** |
| --- | --- | --- | --- | --- | --- | --- |
| Temperature33 / Temperature41 | 0.732677 | 0.583995 | Inf | 1 | -0.39024 | 0.998835 |
| Temperature33 / Temperature42 | 0.20809 | 0.142344 | Inf | 1 | -2.29483 | 0.196013 |
| Temperature33 / Temperature43 | 0.340693 | 0.243043 | Inf | 1 | -1.5094 | 0.658213 |
| Temperature33 / Temperature44 | 0.087734 | 0.057894 | Inf | 1 | -3.68768 | **0.003097** |
| Temperature33 / Temperature45 | 0.081873 | 0.053872 | Inf | 1 | -3.80333 | **0.001981** |
| Temperature41 / Temperature42 | 0.284013 | 0.175427 | Inf | 1 | -2.03787 | 0.320724 |
| Temperature41 / Temperature43 | 0.464997 | 0.302717 | Inf | 1 | -1.17621 | 0.848357 |
| Temperature41 / Temperature44 | 0.119744 | 0.070816 | Inf | 1 | -3.58882 | **0.004485** |
| Temperature41 / Temperature45 | 0.111745 | 0.065892 | Inf | 1 | -3.71658 | **0.002774** |
| Temperature42 / Temperature43 | 1.637239 | 0.826108 | Inf | 1 | 0.977085 | 0.925331 |
| Temperature42 / Temperature44 | 0.421614 | 0.179693 | Inf | 1 | -2.02642 | 0.327137 |
| Temperature42 / Temperature45 | 0.393449 | 0.166808 | Inf | 1 | -2.20019 | 0.237508 |
| Temperature43 / Temperature44 | 0.257515 | 0.121599 | Inf | 1 | -2.8731 | 0.046793 |
| Temperature43 / Temperature45 | 0.240313 | 0.113325 | Inf | 1 | -3.02352 | **0.030065** |
| Temperature44 / Temperature45 | 0.933198 | 0.356807 | Inf | 1 | -0.18083 | 0.999973 |

ST5 Environmental data (n of loggers of each here)

| **Habitat/recorder location** | **Mean Daily Max Temp** | **Max Daily Max Temp** | **SD** | **Top 10% Mean** | **Median** | **90th Percentile** | **IQR** | **CV (%)** |
| --- | --- | --- | --- | --- | --- | --- | --- | --- |
| Snail biologgers on mangroves (emmersed) | 33.97 | 50.5 | 5.6 | 44.51 | 33.5 | 41.5 | 8 | 16.49 |
| Water below mangroves (.25 -.5m depth) | 30.54 | 33 | 1.44 | 32.71 | 30.5 | 32.2 | 2 | 4.71 |
| Water in shallow reef (5m depth) | 30.28 | 31.88 | 0.88 | 31.51 | 30.48 | 31.23 | 1.03 | 2.91 |

ST6 Comparison of model fits for C. sapidus heart rate tpc (n= 12 individuals) in rTPC

| **Model** | **sigma** | **AIC** | **AICc** | **BIC** | **df.residual** |
| --- | --- | --- | --- | --- | --- |
| briere2_1999 | 53.34976 | 210.5493 | 215.1647 | 215.2715 | 15 |
| lactin2_1995 | 40.23487 | 199.8282 | 204.4436 | 204.5504 | 15 |
| gaussian_1987 | 48.00633 | 206.5389 | 211.1543 | 211.2611 | 15 |
| ratkowsky_1983 | 26.96017 | 184.614 | 189.2294 | 189.3362 | 15 |
| quadratic_2008 | 48.35905 | 206.0433 | 208.9005 | 209.8211 | 16 |
| rezende | 35.8363 | 195.4288 | 200.0442 | 200.151 | 15 |
| thomas_2012 | 37.32908 | 196.9796 | 201.595 | 201.7018 | 15 |
| beta_2012 | 45.58878 | 205.2646 | 212.2646 | 210.9312 | 14 |
| **sharpeschoolfull_1981** | 13.19502 | 158.7433 | **168.9251** | 165.3543 | 13 |

ST7 comparison of model fits for L. angulifera heart rate tpc (n=12 individuals) in rTPC

| **model_name** | **sigma** | **AIC** | **AICc** | **BIC** | **df.residual** |
| --- | --- | --- | --- | --- | --- |
| briere2_1999 | 13.11262 | 205.2669 | 208.4247 | 211.3612 | 21 |
| lactin2_1995 | 30.76662 | 247.9096 | 251.0675 | 254.004 | 21 |
| gaussian_1987 | 10.6068 | 194.6629 | 197.8208 | 200.7573 | 21 |
| ratkowsky_1983 | 5.734004 | 163.9088 | 167.0667 | 170.0032 | 21 |
| quadratic_2008 | 14.73959 | 210.2779 | 212.2779 | 215.1534 | 22 |
| rezende | 9.187495 | 187.4803 | 190.6382 | 193.5746 | 21 |
| thomas_2012 | 9.191763 | 187.5035 | 190.6614 | 193.5979 | 21 |
| beta_2012 | 12.71099 | 204.4917 | 209.1584 | 211.805 | 20 |
| **sharpeschoolfull_1981** | 5.251567 | 161.0123 | **167.6006** | 169.5445 | 19 |

ST8 comparison of model fits for L. angulifera righting time TPC

| **Model** | **sigma** | **AIC** | **AICc** | **BIC** | **df.residual** |
| --- | --- | --- | --- | --- | --- |
| briere2_1999 | 0.009927 | -77.8124 | -69.241 | -74.9876 | 9 |
| lactin2_1995 | 0.009791 | -78.1715 | -69.6 | -75.3467 | 9 |
| gaussian_1987 | 0.010434 | -76.5166 | -67.9452 | -73.6919 | 9 |
| **ratkowsky_1983** | **0.009542** | **-78.8402** | **-70.2688** | **-76.0155** | **9** |
| quadratic_2008 | 0.010419 | -77.1847 | -72.1847 | -74.9249 | 10 |
| rezende | 0.009664 | -78.5103 | -69.9389 | -75.6856 | 9 |
| thomas_2012 | 0.009521 | -78.8984 | -70.327 | -76.0737 | 9 |
| beta_2012 | 0.010095 | -76.9084 | -62.9084 | -73.5187 | 8 |
| sharpeschoolfull_1981 | 0.010301 | -76.1195 | -53.7195 | -72.1649 | 7 |

S1 Critical thermal maximum for both species

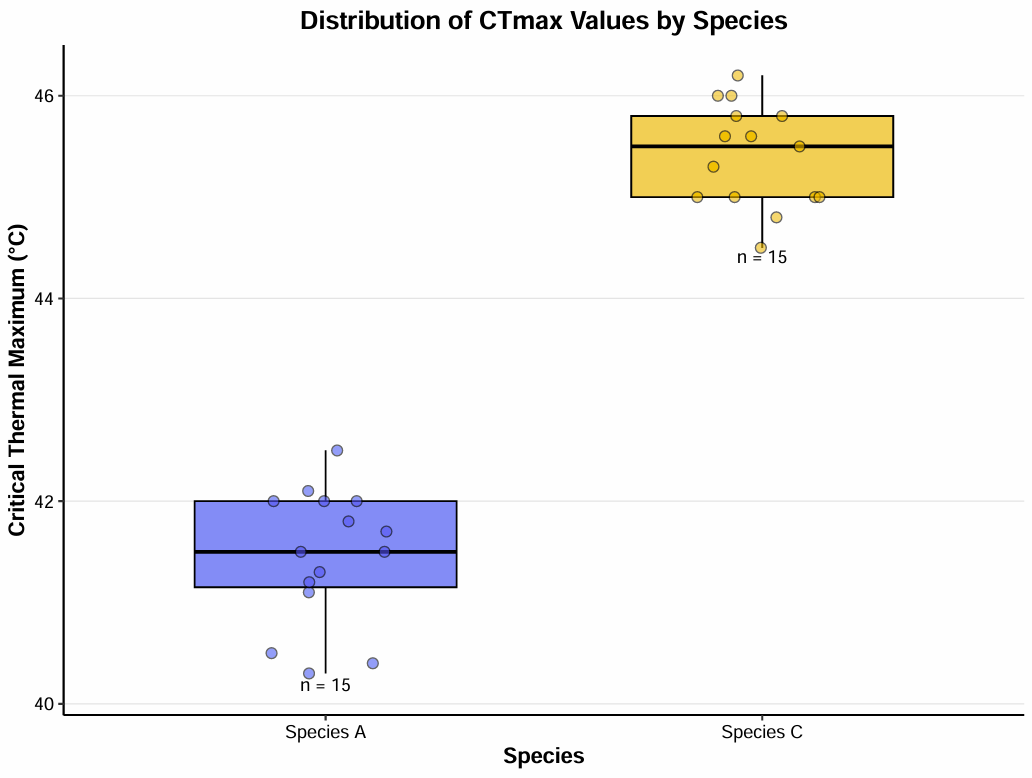

ST9 Summary statistics of Critical thermal maximum in *C. sapidus* and *L. angulifera*

| **Species** | **n** | **Mean (°C)** | **SD** | **SE** | **Median** | **Min** | **Max** |
| --- | --- | --- | --- | --- | --- | --- | --- |
| C. sapidus | 15 | 41.46 | 0.66 | 0.17 | 41.5 | 40.3 | 42.5 |
| L. angulifera | 15 | 45.41 | 0.51 | 0.13 | 45.5 | 44.5 | 46.2 |
